# DUX4 induces a homogeneous sequence of molecular changes, culminating in the activation of a stem-cell-like transcriptional network and induction of apoptosis in somatic cells

**DOI:** 10.1101/2021.05.04.442407

**Authors:** Ator Ashoti, Anna Alemany, Fanny Sage, Niels Geijsen

**Affiliations:** Hubrecht Institute, Developmental Biology and Stem Cell Research, Utrecht, Netherlands; Leiden University Medical Center, Leiden, The Netherlands

## Abstract

Facioscapulohumeral muscular dystrophy (FSHD) is a muscle degenerative disease that disproportionally affects the muscles of the face, shoulder girdle and upper arms. FSHD is caused by the misexpression of Double Homeobox 4 (DUX4), a transcription factor that is normally expressed during early embryonic development. Ectopic expression of DUX4 in somatic cells is cytotoxic and leads to rapid apoptosis. To elucidate the mechanism by which DUX4 induces apoptosis, we determined the temporal transcriptional changes induced by Dux4 at the single-cell level. We observed that induction of DUX4 expression induces a non-random, consecutive sequence of transcriptional changes. DUX4 homogenously induces the activation of a stem cell signature and activates a network of transcription factors that is typically expressed during early embryogenesis and in pluripotent stem cells. Ultimately, these transcriptional changes trigger the induction of apoptosis, suggesting that the induction of this early stemness program is incompatible with a somatic cell program. Our findings shed new light on the timing and dynamic of DUX4-mediated transcriptional reprogramming and may help elucidate why the DUX4 stemness program is required during early embryogenesis, but incompatible with somatic cell viability.

## Introduction

Facioscapulohumeral muscular dystrophy (FSHD) is the third most prevalent muscular dystrophy worldwide^1^. The disease is autosomal dominant, caused by a gain-of-function event, which leads to the ectopic expression of Double Homeobox 4 (DUX4)^2–4^ in affected skeletal muscle, primarily in the face, shoulders and upper arms. DUX4 is a transcription factor normally expressed during early embryonic development^5,6^, in the adult testis^7^ and in the thymus^8^. It induces the expression of a network of genes involved in many different cellular processes, including embryonic development^7,9,10^, RNA processing^9,11,12^, protein homeostasis^12,13^, germline development^7,9,10^, stress response^12,14^, and cell adhesion and migration^11,15^. Expression of DUX4 is stochastic and low, yet potent enough to induce apoptosis in muscle tissue^7,11,16,17^. The mechanism by which DUX4 expression leads to apoptosis downstream of DUX4 is not yet known.

The DUX4 transcription factor is unique to primates and Afrotheria^18,19^ and silenced in most somatic cell types^20^. In FSHD patients DUX4 is stochastically expressed in a burst-like fashion in only 0.1-0.5% of myonuclei^7,17,21^. These aspects have hampered our understanding of the dynamic transcriptional changes that are induced by DUX4. Indeed, in a study of Heuvel et al.^22^, muscle tissue from 4 FSHD patients was analyzed by single-cell RNA sequencing. Out of the 5133 cells that were collected and analyzed from these patients, only 23 cells were classified as DUX4-affected. This reinforces the idea that potential key players might be missed due to the low number of DUX4 affected cells not reaching a critical number needed to detect DUX4-induced transcriptional changes, and/or due to stringency in the analysis.

To study the temporal transcriptional changes that occur upon DUX4 induction, we generated a transgenic cell line, in which DUX4 expression can be induced through the addition of doxycycline^23^. These so called DUX4-inducible expression (DIE) cells allow for precise titration and timing of the DUX4 response. DUX4 induction is robust in DIE cells, as 99-100% of the induced cells enter apoptosis^23^. Using this line, we interrogated whether DUX4 induction led to the induction of defined and orderly molecular changes, or whether it induced a stochastic disruption of gene expression networks before ultimately triggering apoptosis. In order to address this question, we performed single-cell RNA sequencing (SCS) on induced DIE cells, as early as 2 hours after DUX4 induction. By mapping the early molecular changes that follow DUX4 activation at high temporal resolution, we demonstrate that DUX4 induction homogeneously triggers a series of stepwise molecular changes and induces a stem cell-like transcriptional signature with expression of a network of genes that are typically expressed during early embryogenesis. Shortly after the induction of this early embryonic program, the expression of pro-apoptotic genes is detected, suggesting that disruptions caused by the DUX4-induced transcriptional network are incompatible with somatic cell viability.

## Results

### Single cell analysis induced DIE cells

We previously demonstrated that the transcriptomic changes induced by DUX4 in DIE cells are very similar to those reported in FSHD-patient cells and other cellular models^23^. A robust DUX4 expression profile could be seen after only 4.5 hours of induction. However, the manner in which DUX4 expression leads to apoptosis, and what sort of paths are taken is not yet understood. Does DUX4 initiate a defined sequence of transcriptional events every time, or does it initiate a stochastic response that causes a disproportional amount of disruption in the cells that will eventually lead to cell death? To explore this question, we decided to analyze at single-cell resolution the transcriptional changes that occur shortly after DUX4 induction. Our inducible system allows us to examine the immediate effect of DUX4 induction, and track the changes overtime. Due to the robust induction of DUX4, >99% of the DIE cells enter apoptosis withing 48hours of DUX4 induction (data not shown), these changes can be tracked at a high resolution. This type of data would be difficult to attain with primary material, due to the low frequency of DUX4 expression, that occurs in a burst like fashion in a small subset of myonuclei. Furthermore, this inducible system allowed us to time the induction of DUX4, creating a clear timeline trajectory, which is not possible when working with primary material. SCS, will also allow us to detect subtle and perhaps rare early transcriptional changes in specific cell populations that could otherwise be drowned out and missed in bulk RNA sequencing. DIE cells were induced for 2, 3, 4 and 6 hours with doxycycline before sampling and processing for SCS. By reducing the dimensionality using t-Distributed Stochastic Neighbor Embedding (t-SNE) mapping, we were able to have a 2-D visualization of the cell clustering^24,25^. Our results show separate embedding of uninduced cells (0h) and 6h-induced cells, but mixed populations of the intermediate states (2h, 3h and 4h) (Fig. 1A). The cells do however orientate themselves on the y-axis, from the uninduced cells at the top, to the maximum of 6h-induced cells at the bottom (Fig. 1B). This is evident when the expression of known DUX4 target genes were projected onto the t-SNE map (Fig. 1C). LEUTX, PRAMEF1 and ZSCAN4 are genes that have previously shown to increase in expression in FSHD models or FSHD-affected muscle cells^9,11,22,26^. This can indeed also be seen in Fig. 1C, where the expression of these genes is significantly upregulated in 6h-induced cells. This also holds true for genes that are downregulated upon DUX4 expression, such as ID1^9,22^. Figure 1C also shows that as DUX4 induction persists, the expression of ID1 decreases significantly.

**Figure 1.**
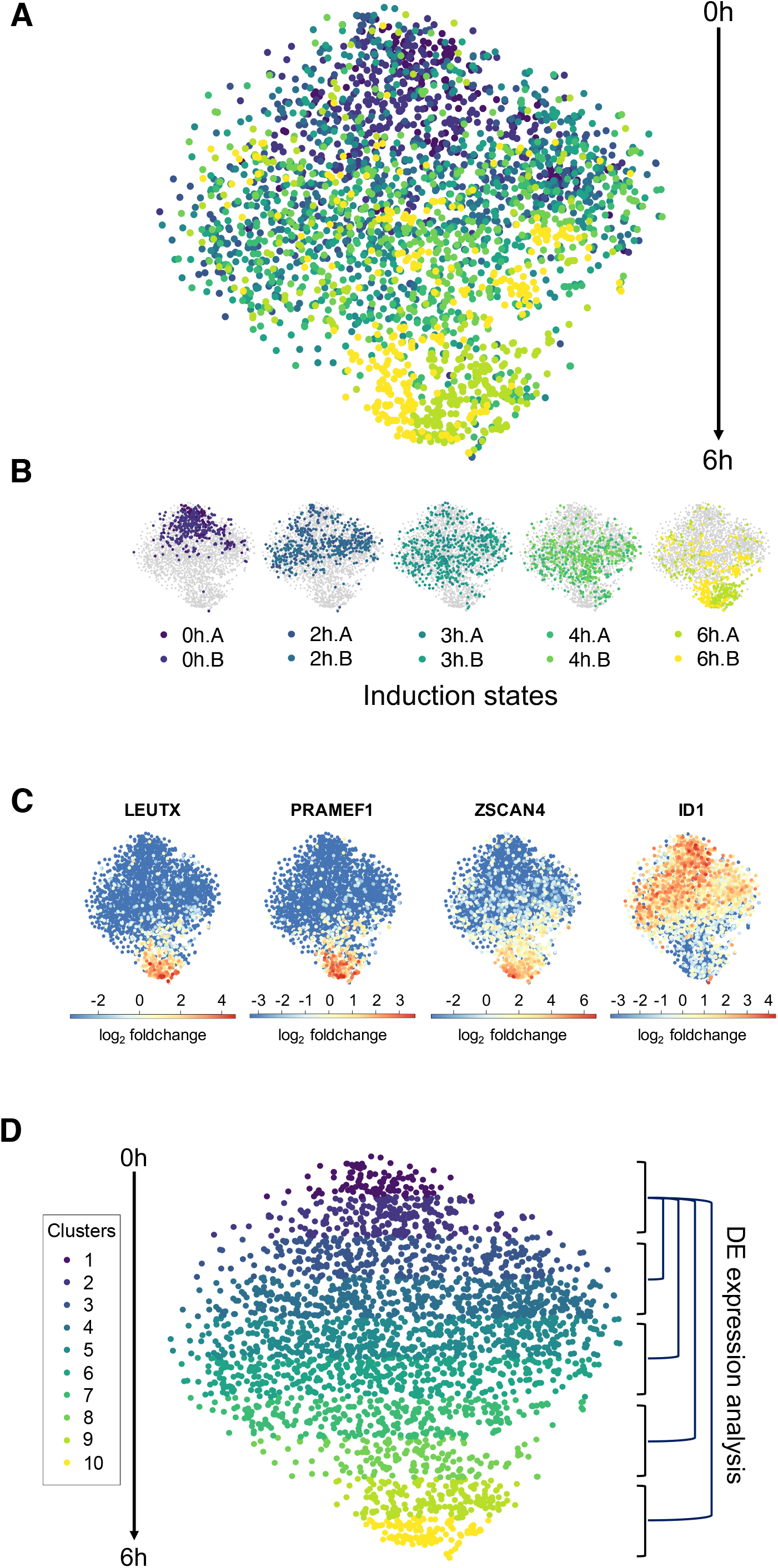
SCS data of DIE cells analyzed with RaceID. **A)** Induced and uninduced DIE single cell data represented in a t-distributed stochastic neighbour embedding (t-SNE) map. Each point represents a single cell, with the induction time of the cells indicated by color. **B)** Individual t-SNE maps for each of the induction time/state. Each sample is indicated in a different color. Induction states are shown from left to right, starting with the two 0h replicates. A and B annotations indicate the two replicates. **C)** t-SNE maps highlighting the expression of DUX4 marker genes. The fold change in gene expression is shown on a log_2_ scale as a linear color scale. **D)** Clustering of the cells in the t-SNE map based on the y-axis coordinates. Clusters are numbered (1-10) and color-coded. Clusters 1 and 2 contain the most uninduced DIE cells, and will be used as the control situation for differential gene expression analysis with the induced clusters (3&4, 5&6, 7&8, and 9&10).

As the cells are organized on the y-axis, we manually divided the vertical axis of the t-SNE into 10 clusters of equal size (Fig. 1D). The mean induction state was calculated for each cluster by considering all cells and their time of induction (Table 1). To avoid confusion, the mean induction state of each cluster will from here on be referred to as the experimental induction times used. Clusters 1 and 2 will therefore be referred to as 0h, 3&4 as 2h, 5&6 as 3h, 7&8 as 4h and 9&10 as 6h. Using this type of clustering, differential gene expression analysis was performed to identify differentially expressed genes between the uninduced cell clusters (clusters 1&2) and the induced cell clusters, as schematically indicated in figure 1D (right).

**Table 1.** Cell make up of t-SNE clusters.

| cluster | 0h.A | 0h.B | 2h.A | 2h.B | 3h.A | 3h.B | 4h.A | 4h.B | 6h.A | 6h.B | cells/<br>cluster | mean<br>state |  |
| --- | --- | --- | --- | --- | --- | --- | --- | --- | --- | --- | --- | --- | --- |
| cl.1 | 33 | 23 | 15 | 10 | 10 | 5 | 0 | 4 | 5 | 1 | 106 | 1.39 | 0h |
| cl.2 | 61 | 77 | 22 | 17 | 25 | 14 | 7 | 11 | 6 | 1 | 241 | 1.28 |  |
| cl.3 | 38 | 106 | 48 | 29 | 30 | 23 | 9 | 17 | 11 | 1 | 312 | 1.57 | 2h |
| cl.4 | 34 | 57 | 83 | 71 | 69 | 58 | 38 | 49 | 18 | 29 | 506 | 2.61 |  |
| cl.5 | 10 | 20 | 54 | 108 | 58 | 51 | 69 | 90 | 15 | 40 | 515 | 3.14 | 3h |
| cl.6 | 1 | 3 | 33 | 41 | 58 | 65 | 64 | 69 | 23 | 48 | 405 | 3.64 |  |
| cl.7 | 0 | 0 | 12 | 14 | 33 | 41 | 76 | 37 | 31 | 39 | 283 | 4.05 | 4h |
| cl.8 | 0 | 0 | 2 | 1 | 15 | 14 | 20 | 11 | 52 | 42 | 157 | 4.97 |  |
| cl.9 | 0 | 1 | 0 | 0 | 2 | 6 | 25 | 5 | 84 | 61 | 184 | 5.51 | 6h |
| cl.10 | 1 | 0 | 1 | 1 | 1 | 2 | 3 | 1 | 71 | 33 | 114 | 5.73 |  |

### Differential gene expression analysis

In order to detect subtle but significant changes in expression, differentially expressed genes were filtered for an adjusted p-value (Padj) of < 10**−6, and a log_2_(foldchange) (log2FC) of > 0.5 and < −0.5 (Tables S1-4). This analysis demonstrated that as the induction time increases, so does the number of differentially expressed genes, with a core group of differentially expressed genes being shared between induction states, (Fig. 2A, Table 2 and Tables S1-4). This suggests that a deterministic chain of events is induced early on, if not immediately after DUX4 expression. Interestingly, even though DUX4 expression itself was not detected in the induced DIE cells, a DUX4 expression profile is readily detected only 3h post DUX4 induction, as can be seen when expression data is entered through the Enrichr database^27,28^ (Fig. 2B). This DUX4 profile even becomes more apparent as induction time increases. The Enricher database can match the entered lists of genes with previously entered studies, matching our gene lists with one other study from Geng et al.^9^ in which DUX4 had been overexpressed in human primary myoblasts. Such an early induced DUX4 expression profile has (to the best of our knowledge) not been seen before, with other studies measuring the effects of DUX4 6h^29^ or 14h^26^ post induction, or 24-36h post lentiviral transfection^9,26^. As no other study has examined the effects of DUX4 at such early time points, our datasets uniquely show the earliest DUX4 affected genes. Furthermore, previously identified DUX4 affected genes such as RFPL4B, GOLGB1, ZNF296, SRSF8, ID1 and ID3^9,11,22,26^, could too be classified as potential early marker genes, as they have been identified as being differentially expressed after a mere 3h of DUX4 induction.

**Table 2:** Summary of differentially expressed genes found in induced DIE cells at different induction times/states.

| Induction state | 2h | 3h | 4h | 6h |
| --- | --- | --- | --- | --- |
| Upregulated genes | 43 | 77 | 133 | 248 |
| Downregulated genes | 8 | 36 | 40 | 68 |
| Differentially expressed genes<br>total | 51 | 113 | 173 | 361 |
| Transcription- and co-factors | 12 | 28 | 44 | 81 |
| Kinases | 3 | 3 | 6 | 10 |

**Figure 2.**
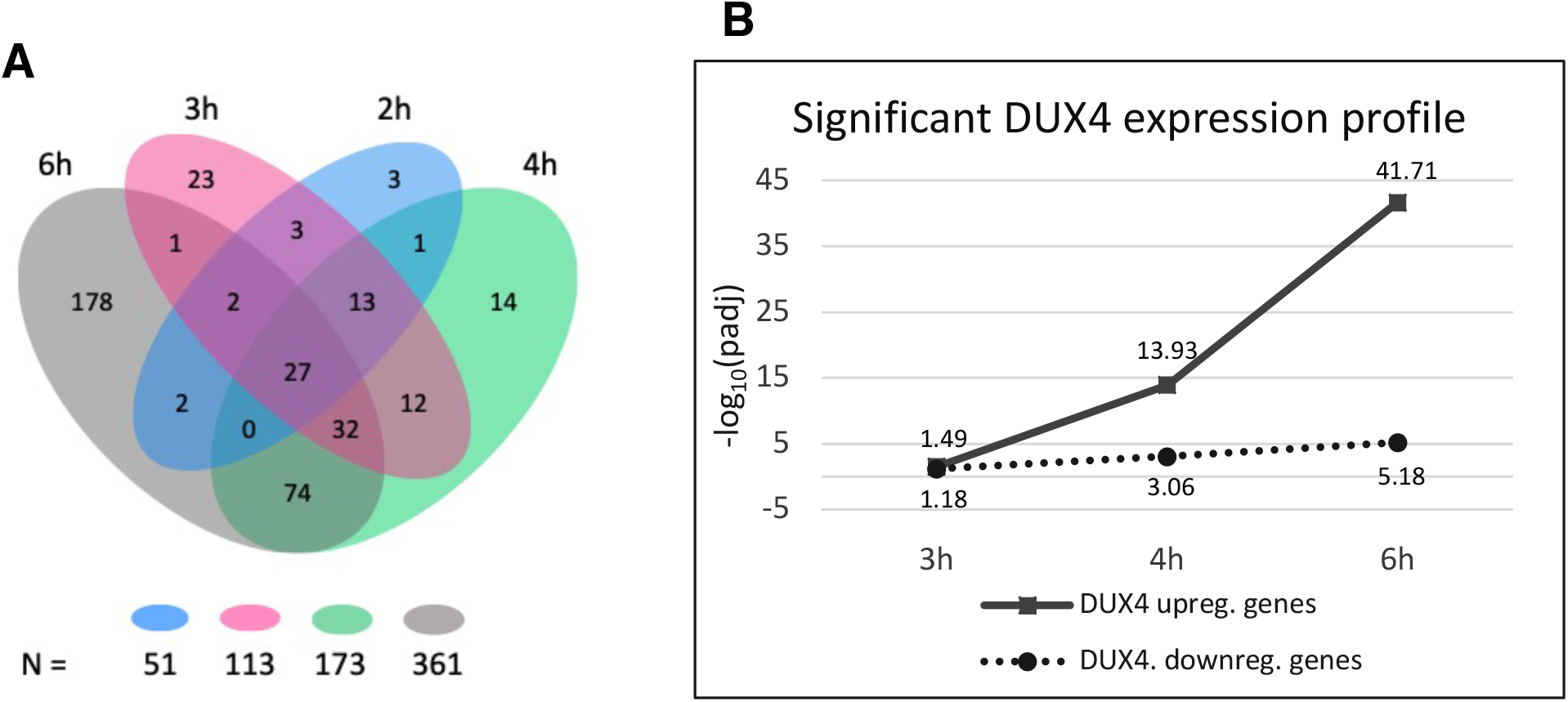
DUX4 differential gene expression profile between induction states. **A)** Venn diagram demonstrating the number of shared differentially expressed genes between different induction times. Total number of differentially expressed genes in each state is shown below the graph, in the same color-coding. **B)** A graph demonstrating the increase in significance of the upregulated and downregulated expression profile of DUX4 in induced DIE cells, according to entries in the Enrichr database. Induction samples are plotted on the x-axis, and the y-axis displays the negative log_10_ of the adjusted p-value (padj), of the detected DUX4 expression profile.

Remarkably, a high percentage (29-36%) of upregulated transcripts encode for transcription factors and cofactors, in addition to a number of differentially expressed kinases (Table 2, Table S5), which was also observed in previous bulk-seq experiments^23^. This suggests that DUX4 induces a network of downstream transcription factors that in turn induces a cascade of secondary transcriptional events, ultimately leading to apoptosis. Since the expression of transcription factors can be low, many additional factors might fall under the detection limit in single-cell sequencing data. Enrichr^27,28^ was therefore used to analyze our datasets for the presence of signature gene expression profiles, or “transcriptional footprints”, that are indicative of the activity of specific transcription factors. Analysis of our DIE cell data using Enrichr yielded a list of potential transcription factors that can explain the observed changes in gene expression (Table S6). Some of the identified profiles did indeed match differentially expressed transcription factors in the induced DIE cells (SOX3, NR2F2, ZNF217 and OTX2). In addition, the Enrichr algorithm detected several other transcription factor profiles of transcription factors which themselves are not found to be differentially expressed in our dataset (Table S6). A number of these “transcriptional footprints” were found in all 4 timepoints examined (LIN28, SOX5, ZIC3, JUNB, KLF10, MEIS2, MYCN, PITX, SETB1, ZEB2, ZNF503, MYB, WT1, NR2F2), suggesting that these factors are induced early after DUX4 induction and persist with continued DUX4 expression. Of particular interest is the identification of several transcription factors which are represented in both the upregulated as well as the downregulated gene set (e.g. ZIC3, JUNB, KLF10, MEIS2, MYCN and SETB1). As both upregulated and downregulated genes corresponded to the activity status of these transcription factors, it does strongly suggest their role in the DUX4-induced cytotoxic cascade, as opposed to only finding a one-sided effect.

In summary, we have found that the induction of DUX4 promotes changes in the transcriptional landscape of DIE cells as early as 2 hours post doxycycline administration. Many differentially expressed genes found in the early data sets maintain their differentially expressed status with continued DUX4 expression. This corroborates the notion that DUX4 initiates a clear progressive cascade of events, and does not stochastically and/or randomly affects genes and pathways. This is further corroborated by our finding that transcription factors and co-factors are overrepresented in the list of differentially expressed genes and account for approximately ~33% of the differentially upregulated genes, whereas they only comprise 11-13.5% of the human genome. This indeed suggests that DUX4 activates a coherent network of transcriptional regulators that together initiate a new cellular program that ultimately leads to cell death. Lastly the presence of transcription factors could also be deduced from the detection of their transcriptional footprint, even when the expression of the individual factors themselves were not always detected. This is a common shortcoming of single cell sequencing, where the detection of low abundant transcripts can be missed.

### Gene ontology

To identify which biological processes are affected by the temporal changes in gene expression in the induced DIE cells, the differentially upregulated and downregulated genes were analyzed using the PANTHER (<u>P</u>rotein <u>AN</u>alysis <u>TH</u>rough <u>E</u>volutionary <u>R</u>elationships) algorithm. PANTHER is an online tool that classifies proteins (and their corresponding genes) based on their family or subfamily, their molecular function, and their involvement in any biological processes and pathways, to facilitate high throughput analysis of datasets^30–32^. The biological processes that were identified are shown in Fig. 3A and Table S7. Gene ontology (GO) terms were assigned a general “umbrella” term. Table S7 shows the full list of GO terms.

**Figure 3.**
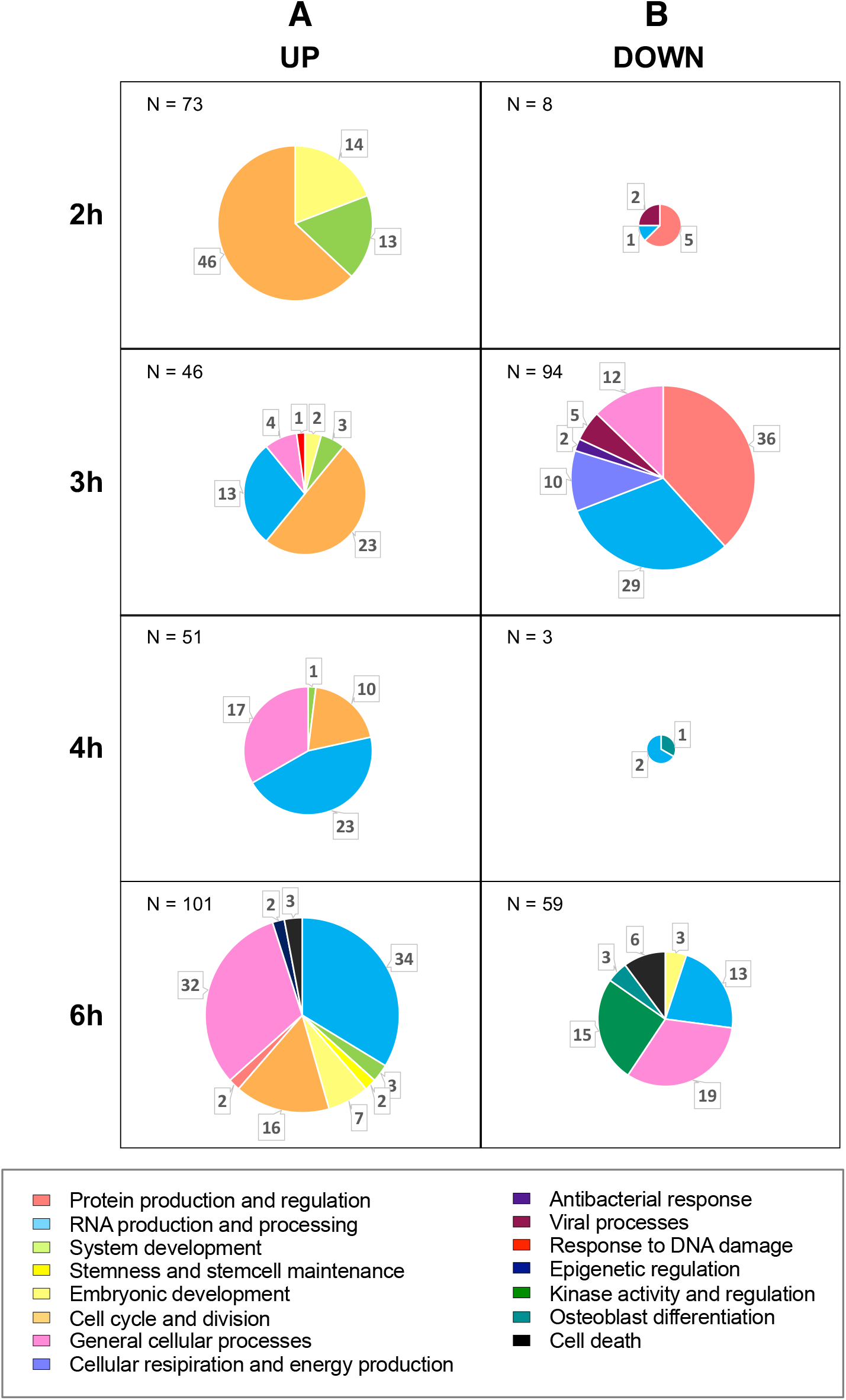
Gene ontology reveals DUX4-induced paths. Gene ontology performed on gene lists of differentially **A)** upregulated and **B)** downregulated genes from the 4 induction states. Only biological processes with an FDR < 0.05, and a raw p value of < 10**−3 are included. Biological processes are color-coded based on their general “umbrella” term. The total number of detected biological processes is indicated in the top left corner of each diagram, with the number of biological processes per umbrella term annotated in the pie chart. See table S7 for the full list of biological processes per induction state.

As shown, DUX4-induction initially triggered the activation of an early developmental program and processes involved in the cell cycle and proliferation. Three hours after induction, genes involved in developmental processes are less prominent, with the majority of processes now being involved in cell cycle and RNA processing. At 6h of induction, the first apoptotic processes were identified. GO terms identified with the downregulated genes appeared more incoherent than the GO terms detected with the upregulated gene sets. This could be due to the nature of DUX4 being more of a transcriptional activator, rather than a repressor^10^. GO terms found with the downregulated gene stets might thus reflect the loss of cell identity, consistent with the idea that DUX4 initiates an early embryonic transcriptional program. Nonetheless, processes involved in programmed cell death were also found upon analyzing the downregulated genes after 6h of DUX4 induction (Fig. 3B). Furthermore, at 3h post DUX4-induction, downregulated genes demonstrate changes in cellular respiration and energy production. These processes contribute to oxidative stress, a common occurrence in FSHD-affected cells that is likely involved in DUX4-induced apoptosis^33–35^.

The temporal identification of altered biological processes revealed a sequential path that is activated upon DUX4 induction. This path starts by activating developmental processes, and subsequently many other processes involved in RNA processing, protein production and regulation, cellular respiration, kinase activity, eventually leading to the induction of apoptosis.

### StemID

The observation that DUX4 initially activates an early embryonic transcriptional program was interesting and suggests that DUX4 temporarily converts cells toward a developmentally immature state. This notion is corroborated by StemID, an algorithm that uses transcriptome entropy to identify stem cells within a cell population^36^. Determining transcriptome diversity in single cells is done by using Shannon's entropy^37^, which measures disorder in high-dimensional systems. The entropy value of a given cell type indicates the degree of transcriptomic promiscuity. As pluripotent stem cells have the option of differentiation in any cell type, a wide number of signaling pathways need to remain active, which is reflected as high transcriptome entropy. As these cells become more committed to a specific cell fate, the number of active pathways decrease to a few specific pathways needed to maintain their cell identity, which in turn leads to a decrease in transcriptome entropy^36,38,39^. When a lineage trajectory is projected onto the t-SNE map, it becomes clear by the color indication of the vertices that transcriptome entropy peaks in cluster 4 (Fig. 4A). The barplot in Fig. 4B clearly shows the increase in transcriptome entropy, until it peaks in cluster 4, which has an induction state of around 2h. Transcriptome entropy then slowly decreased with increasing induction times. This is thus corroborating gene ontology results that showed the induction of a more embryonic developmental state in ~2h induced DIE cells (clusters 3&4).

**Figure 4.**
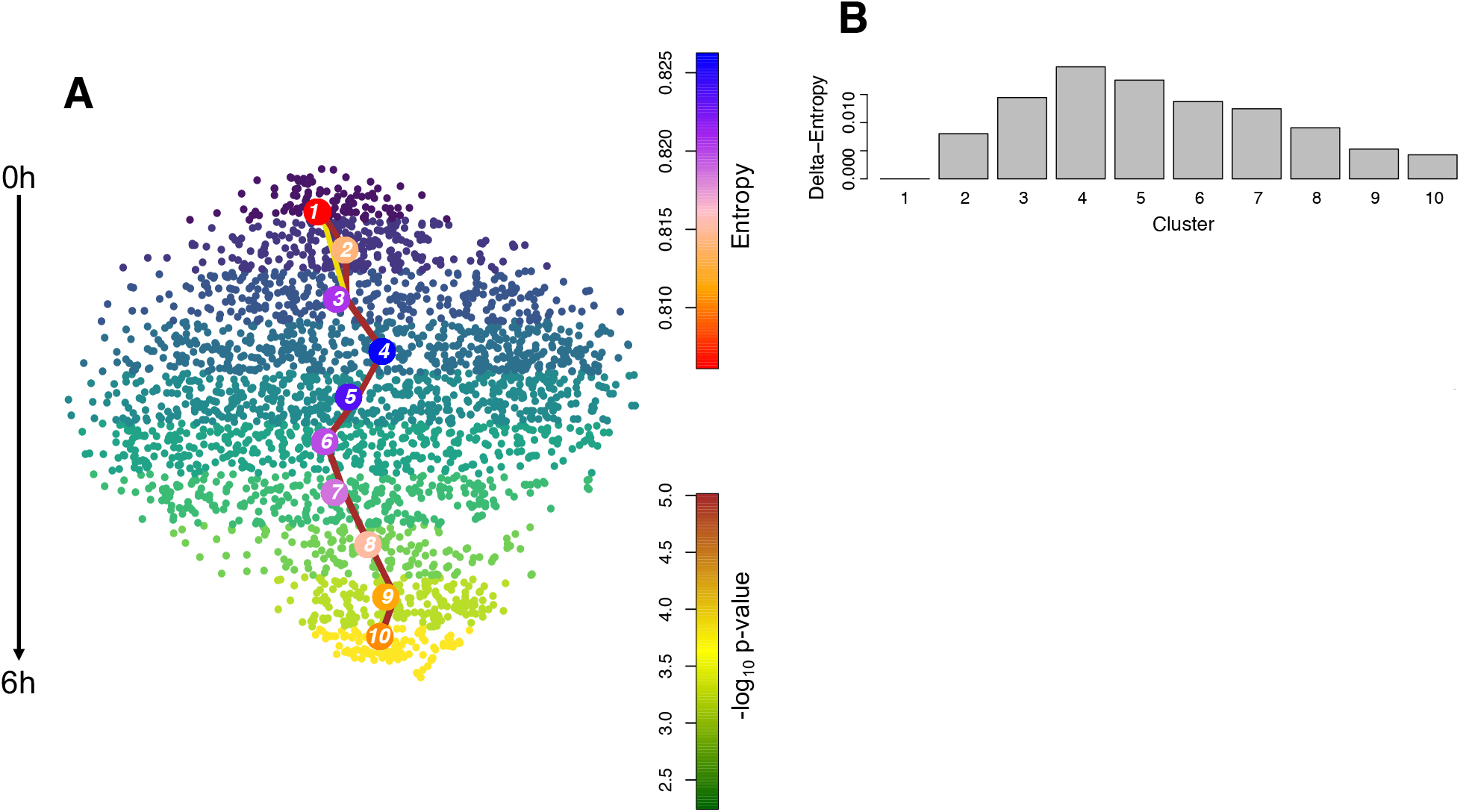
StemID analysis identified a stem cell state in single cell clusters. **A)** Inferred lineage tree superimposed onto the t-SNE map. The level of entropy of a cluster is indicated by the color of the vertices. The link color between vertices represents the −log_10_ value, with only the significant links being shown (p < 0.01). **B)** Barplot showing the delta-entropy score per cluster. The delta entropy was calculated by subtracting the lowest entropy score across all cells from the median transcriptome entropy of each cluster.

### Pseudotime analysis with FateID

StemID revealed a transient trajectory of stemness in induced DIE clusters, which peaked at cluster 4 and was subsequently followed by a gradual decrease of stemness at later time points, when further transcriptome changes reflect profound changes in metabolism and RNA processing. To follow up on this observation, pseudotime analysis was employed to further define temporal stages of transcriptional states, using FateID^40^. By doing so, we were able to identify stage-specific co-expression patterns across this vertical trajectory based on previous t-SNE clustering (Fig. 1D). Expression patterns of known DUX4 target genes show a gradual increase (LEUTX, ZSCAN4, ZNF217, and PRAMFE1), or decrease in expression (ID1 and ID3) as DUX4 expression persisted (Fig. 5A), which is in line with earlier observations seen above (Fig. 1A) and previous observations in other studies^9,11,22,26^. These genes were present in gene nodes that showed a gradual increase or decrease in gene expression (e.g. nodes 18-21 or 1-3 respectively) (Fig. 5B). Moreover, dynamic gene expression patterns were identified in other gene nodes, such as oscillating expression patterns during the 6 hours of DUX4 induction (e.g. 4, 6, 10 and 17) (Fig. 5B and S2). This suggests the activation of a very dynamic underlying process, upon DUX4 induction, in which some genes are induced and inhibited multiple times in a relatively short time frame.

**Figure 5.**
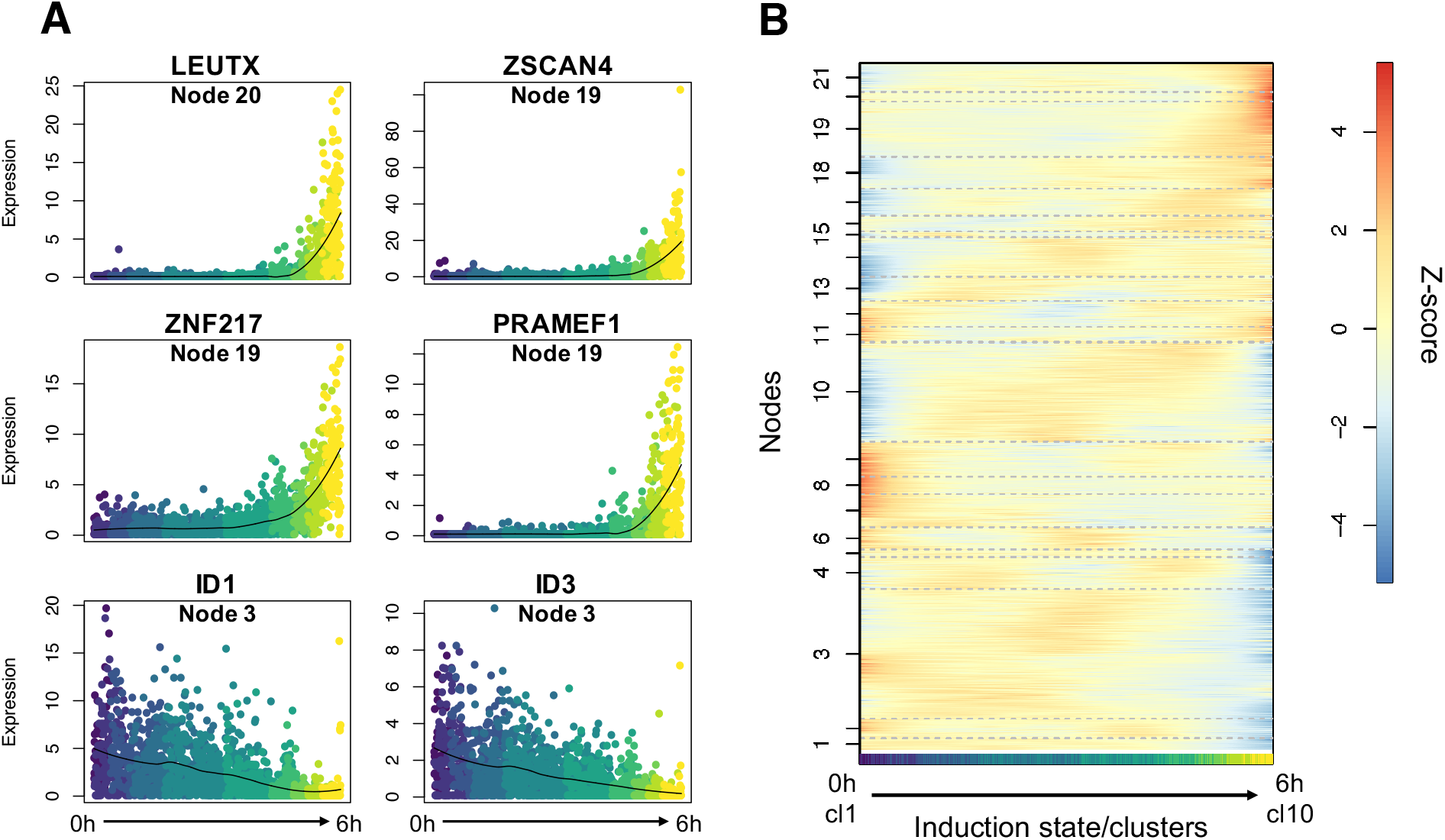
Gene expression patterns in the induced DIE cell trajectory by FateID pseudotime analysis. **A)** Expression patterns of 6 known DUX4 marker genes following the DIE cell induced trajectory (0h-6h induction). The nodes in which these genes are contained are annotated below the gene name in brackets. **B)** Self-organizing heatmap of z-score transformed pseudotime expression profiles across the DIE cell induced trajectory (0h-6h induction), based on the t-SNE map. Cells are represented on the x-axis, and the genes are organized in nodes that are represented on the y-axis. Genes with a similar expression pattern are clustered in nodes, with a color indication representing gene expression, based on their transformed z-score.

Analysis of the differentially expressed genes in the oscillating nodes did not yield a clear answer as to why these particular sets of genes vary in their expression during DUX4 induction. Of the nodes that demonstrated clear oscillating patterns (4, 6, 10, 12, 17), node 4 and 6 did not contain differentially expressed genes, node 12 contained one differentially downregulated gene (COX7A2), node 10 contained 27 differentially upregulated genes, and node 17 contained 17 differentially upregulated genes (Table 3). Using the STRING database^41,42^, we were able to determine that the differentially expressed genes from node 10 are primarily involved in developmental process, system development, and cell cycle and division, with many genes interconnecting and involved in all three processes. STRING is a database of known and predicted protein-protein interactions (both direct and indirect), that allowed us to visualize the types of associations between genes (Fig. 6A). The biological processes identified with STRING are similar to those identified using the PANTHER algorithm (Fig. 3A). The expression pattern in node 10 therefore fits previous GO analyses, demonstrating a temporal increase in the number of developmental genes and processes, peaking at around 2-3h (Fig. 5B and 6B). Differentially upregulated genes in node 17 did not show to be part of any significantly affected biological processes, nor a clear coherent core network could be seen between the genes as was found in node 10.

**Figure 6.**
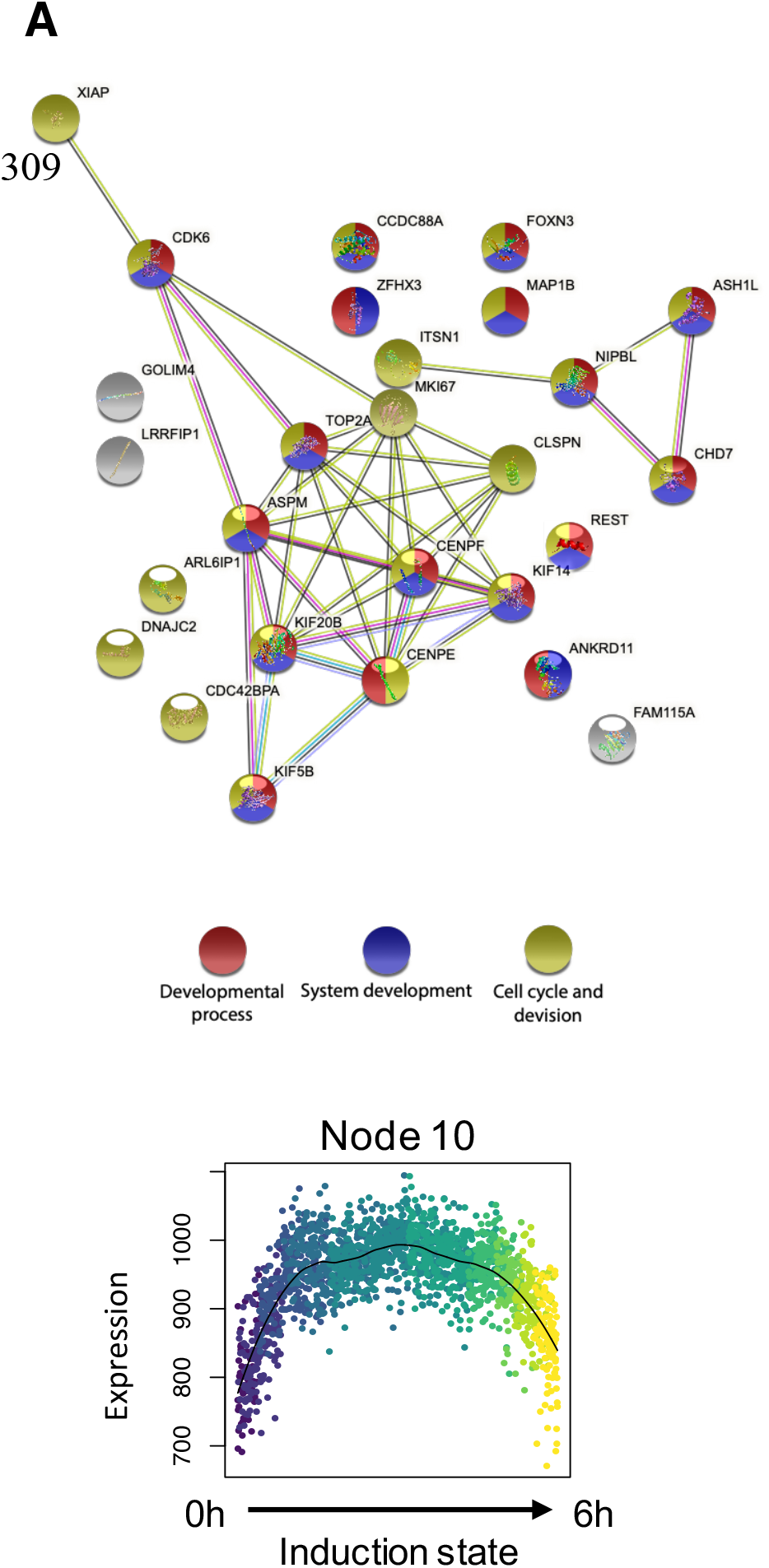
Schematic representation of gene affiliations within node 10 using STRING^41,42^. **A)** Affiliations between differentially upregulated genes of node 10. Genes involved in the developmental process are shown in red, in system development in blue, and in the cell cycle in yellow. Genes indicated in white are involved in other general molecular processes. The color of the links indicates the nature of the affliation between genes. Pink and cyan represent known interactions that were experimentally determined, or from curated databases respectively. Green, red and blue interactions represent predicted interactions based on gene neighborhood, gene fusions and gene occurrence respectively. Yellow, black and purple interactions are based on textmining, co-expression, and protein homology respectively. Images were adapted from STRING-derived interaction networks. **B)** Expression pattern graph of node 10 genes, following the DIE cell induced time trajectory (0h-6h induction).

## Discussion

To interrogate the temporal transcriptional changes that are induced by the transcription factor DUX4, we applied a novel cell model in which DUX4 expression can robustly be induced. Our data reveals at the single-cell level, how DUX4 induction lead to the activation of a transient early embryonic and stemness state, by activating a network of early embryonic genes, of which a disproportionally large fraction are themselves transcription factors. Our analysis suggests that DUX4 initiates a non-random consecutive chain of events that is exacerbated as time progresses. In addition, analysis of gene expression profiles using Enrichr revealed that transcriptional signatures of transcription factors, most of which were not detected during the induction periods, are already prevalent 2 hours post-induction and indeed became more profound as induction time progresses. As such, we have uncovered a larger network of transcription factors that might play a role in triggering subsequent molecular changes that ultimately contribute to DUX4 -mediated cytotoxicity.

Our results suggest an initial activation of developmental processes that lead cells to an increased stemness state only 2 hours after DUX4 induction. Next, additional biological processes such as RNA processing, and protein production and regulation were activated, eventually leading to the activation of apoptotic processes 6 hours post DUX4 induction.

The analysis performed in this study revealed which genes started diverging in their expression during the first few hours of DUX4 induction. At these early timepoints, some transcriptional changes were subtle, but as these changes in expression remained or even intensified with increased induction times, we believe them to be significant. More attention should be focused towards some of these subtle changes in expression at these early post-induction timepoints, that could normally be missed due to too stringent filtering. DUX4 itself proves that the smallest changes in expression can cause major consequences. This factor was not identified in our transcriptomic analysis, concurrent with its low transcriptional and abundance levels in muscle tissue of FSHD patients^3,7,17,21^. While key transcription factors, such as DUX4, are themselves not detected, they leave behind a detectable “footprint” in the transcriptome of affected cells. These elusive genes could have great implications in our understanding of the role of DUX4 in early embryogenesis, as well as in the pathophysiology of FSHD.

## Methods

### Cell culturing and seeding

DIE cells were cultured in growth medium consisting of IMDM basal medium with 10% Tet system-approved FBS (Clontech) and 55μM 2-mercaptoethanol, supplemented with 5μg/ml Puromycin and 6μg/ml Blasticidin.

### Sample preparation and SORT-seq

DIE cells were grown in 48-wells plates, until a ~90% confluency was reached. Cells were exposed to 1μg/ml doxycycline for 2, 3, 4, and 6 hours. Doxycycline-exposed and untreated DIE cells were rinsed with DPBS after which 0.25% Trypsin-EDTA (Thermo Scientific) was added. Trypsin was immediately removed after it had covered the complete surface. Cell were incubated for 1 minute at 5% CO_2_ and 37°C, after which the trypsin was deactivated by adding IMDM media supplemented with 10% Tet-system approved FBS and DAPI nuclear stain. Trypsinized DIE cells were resuspended in the media and then strained using Cell-strainer capped tubes (Falcon). Cells were stored on ice until FACS sorting. Viable DAPI negative cells were sorted into 384 hard shell plates (Biorad) with 5 μl of vapor-lock (QIAGEN) containing 100-200 nl RT primers, dNTPs, and synthetic mRNA Spike-Ins, using the FACSJazz (BD biosciences). The plates were immediately spun down and stored a −80°C. Cells were processed as described in Muraro et al.^43^, using the CEL-seq2-bases scRNA-seq. Samples were sequenced using Illumina Nextseq 500, 2×75 kit, high output. Two biological replicates per samples were sent for sequencing. Initial normalization and mapping were done as described by Muraro et al^43^.

## Data analysis

Illumina sequencing-generated paired-end reads were aligned, mapped, and normalized as previously described^43^. For single cell analysis, cells with a minimum of 6000 transcripts were considered, and data normalization was performed by downsampling transcript counts to 6000 for all cells (Fig. S1). Initial analysis revealed a batch affect between the two uninduced biological replicates. One sample showed signs of additional metabolic stress. The top 187 diverging genes (padj < 10**−7) between the two uninduced biological replicates were removed from all data to account for any source of metabolic stress. Dimensionality reduction of cells was done using RaceID^25^, after which the clusters were manually determined by dividing the y-axis of the tSNE map in 10 clusters of equal size. Differential expression of genes between cell clusters were identified as described by Muraro et al^43^, based on a previous publication of Anders and Huber^44^. Pseudotime analysis was performed using StemID^36^ and FateID^40^.

## Data Resources

The RNA sequencing datasets are available from the GEO data base, accession number: GSE156154.

## Acknowledgements

This study was supported by Stichting FSHD and the SingelSwim Utrecht.

## Supplementary data

**Figure S1.**
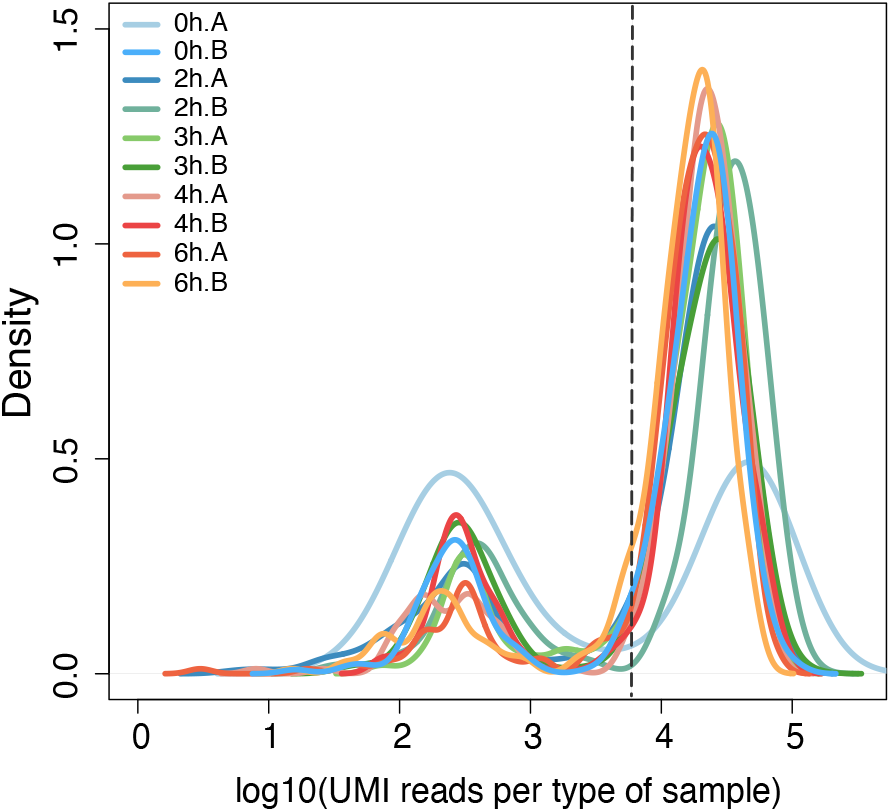
Density plot representing the total read count of all samples. Samples are color coded. An A or B annotation represents to which biological replicate the sample belongs. The intermittened line shows the cutoff of the number of UMI reads (6000) used to determine which cells to inlcude for the analysis. All cells with a normalized transript count above 6000 have been included in the RaceID analysis.

**Figure S2.**
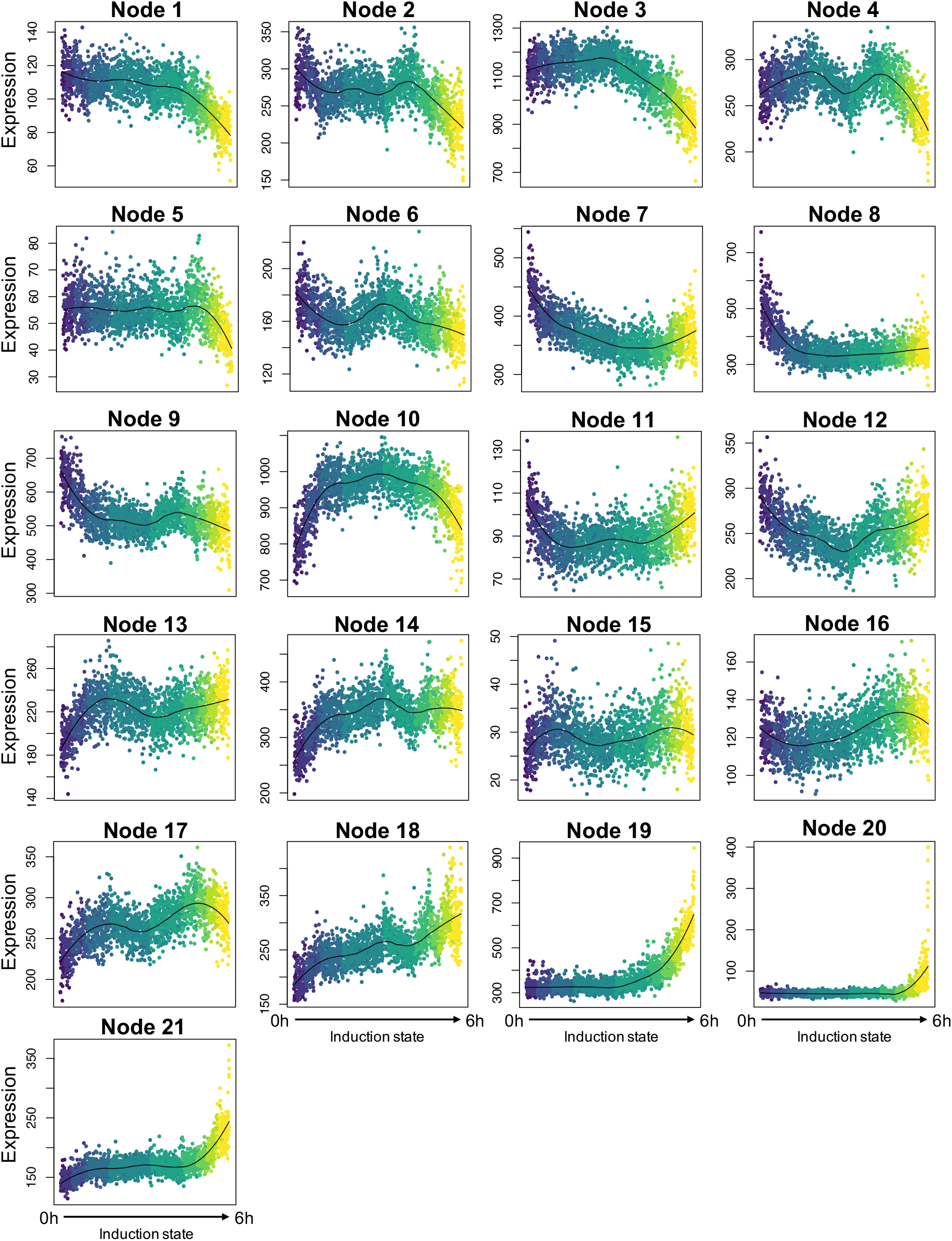
Gene expression patterns of gene nodes from pseudotime analysis. Dynamic gene expression patterns of all nodes of the self-organizing heat map of figure 5A. Each point represents a cell. The color of the point and its location on the x-axis represents its induction state from uninduced (0h) to 6h induced. Normalized expression is plotted on the y-axis. The black line indicates a local regression.

## Notes

### Competing Interest Statement

The authors have declared no competing interest.

## References

1. Deenen, J. C. W. et al. Population-based incidence and prevalence of facioscapulohumeral dystrophy. Neurology 83, 1056–1059 (2014).

2. Kowaljow, V. et al. The DUX4 gene at the FSHD1A locus encodes a pro-apoptotic protein. Neuromuscul. Disord. 17, 611–623 (2007).

3. Dixit, M. et al. DUX4, a candidate gene of facioscapulohumeral muscular dystrophy, encodes a transcriptional activator of PITX1. Proc. Natl. Acad. Sci. U. S. A. 104, 18157–18162 (2007).

4. Snider, L. et al. RNA transcripts, miRNA-sized fragments and proteins produced from D4Z4 units: New candidates for the pathophysiology of facioscapulohumeral dystrophy. Hum. Mol. Genet. 18, 2414–2430 (2009).

5. Whiddon, J. L., Langford, A. T., Wong, C. J., Zhong, J. W. & Tapscott, S. J. Conservation and innovation in the DUX4-family gene network. Nat. Genet. 49, 935–940 (2017).

6. Hendrickson, P. G. et al. Conserved roles of mouse DUX and human DUX4 in activating cleavage-stage genes and MERVL/HERVL retrotransposons. Nat. Genet. 49, 925–934 (2017).

7. Snider, L. et al. Facioscapulohumeral dystrophy: Incomplete suppression of a retrotransposed gene. PLoS Genet. 6, 1–14 (2010).

8. Das, S. & Chadwick, B. P Influence of repressive histone and DNA methylation upon D4Z4 transcription in non-myogenic cells. PLoS One 11, 1–26 (2016).

9. Geng, L. N. et al. DUX4 Activates Germline Genes, Retroelements, and Immune Mediators: Implications for Facioscapulohumeral Dystrophy. Dev. Cell 22, 38–51 (2012).

10. Knopp, P. et al. DUX4 induces a transcriptome more characteristic of a less-differentiated cell state and inhibits myogenesis. J. Cell Sci. 129, 3816–3831 (2016).

11. Rickard, A. M., Petek, L. M. & Miller, D. G. Endogenous DUX4 expression in FSHD myotubes is sufficient to cause cell death and disrupts RNA splicing and cell migration pathways. Hum. Mol. Genet. 24, 5901–5914 (2015).

12. Jagannathan, S., Ogata, Y., Gafken, P. R., Tapscott, S. J. & Bradley, R. K. Quantitative proteomics reveals key roles for post-transcriptional gene regulation in the molecular pathology of facioscapulohumeral muscular dystrophy. Elife 8, 1–16 (2019).

13. Homma, S., Beermann, M. Lou, Boyce, F. M. & Miller, J. B. Expression of FSHD-related DUX4-FL alters proteostasis and induces TDP-43 aggregation. Ann. Clin. Transl. Neurol. 2, 151–166 (2015).

14. Winokur, S. T. et al. Facioscapulohumeral muscular dystrophy (FSHD) myoblasts demonstrate increased susceptibility to oxidative stress. Neuromuscul. Disord. 13, 322–333 (2003).

15. Bosnakovski, D. et al. Transcriptional and cytopathological hallmarks of FSHD in chronic DUX4-expressing mice. J. Clin. Invest. 130, 2465–2477 (2020).

16. Bosnakovski, D. et al. Muscle pathology from stochastic low level DUX4 expression in an FSHD mouse model. Nat. Commun. 8, 1–9 (2017).

17. Tassin, A. et al. DUX4 expression in FSHD muscle cells: How could such a rare protein cause a myopathy? J. Cell. Mol. Med. 17, 76–89 (2013).

18. Clapp, J. et al. Evolutionary conservation of a coding function for D4Z4, the tandem DNA repeat mutated in facioscapulohumeral muscular dystrophy. Am. J. Hum. Genet. 81, 264–279 (2007).

19. Leidenroth, A. et al. Evolution of DUX gene macrosatellites in placental mammals. Chromosoma 121, 489–497 (2012).

20. Van der Maarel, S. M., Tawil, R. & Tapscott, S. J. Facioscapulohumeral muscular dystrophy and DUX4: Breaking the silence. Trends Mol. Med. 17, 252–258 (2011).

21. Jones, T. I. et al. Facioscapulohumeral muscular dystrophy family studies of DUX4 expression: Evidence for disease modifiers and a quantitative model of pathogenesis. Hum. Mol. Genet. 21, 4419–4430 (2012).

22. Van Den Heuvel, A. et al. Single-cell RNA sequencing in facioscapulohumeral muscular dystrophy disease etiology and development. Hum. Mol. Genet. 28, 1064–1075 (2019).

23. Ashoti, A. et al. A genome-wide CRISPR/Cas phenotypic screen for modulators of DUX4 cytotoxicity reveals screen complications. bioRxiv 2020.07.27.223420 (2020) doi:10.1101/2020.07.27.223420.

24. van der Maaten, L. & Hinton, G. H. Visualizing Data using t-SNE. J. Mach. Learn. Res. 9, 2579–2605 (2008).

25. Grün, D. et al. Single-cell messenger RNA sequencing reveals rare intestinal cell types. Nature 525, 251–255 (2015).

26. Jagannathan, S. et al. Model systems of DUX4 expression recapitulate the transcriptional profile of FSHD cells. Hum. Mol. Genet. 25, ddw271 (2016).

27. Chen, E. Y. et al. Enrichr: Interactive and collaborative HTML5 gene list enrichment analysis tool. BMC Bioinformatics 14, (2013).

28. Kuleshov, M. V. et al. Enrichr: a comprehensive gene set enrichment analysis web server 2016 update. Nucleic Acids Res. 44, W90–W97 (2016).

29. Choi, S. H. et al. DUX4 recruits p300/CBP through its C-terminus and induces global H3K27 acetylation changes. Nucleic Acids Res. 44, 5161–5173 (2016).

30. Ashburner, M. et al. Gene Ontology: tool for the unification of biology. Nat. Genet. 25, 25–29 (2000).

31. Carbon, S. et al. The Gene Ontology Resource: 20 years and still Going strong. Nucleic Acids Res. 47, D330–D338 (2019).

32. Mi, H., Muruganujan, A., Ebert, D., Huang, X. & Thomas, P. D. PANTHER version 14: More genomes, a new PANTHER GO-slim and improvements in enrichment analysis tools. Nucleic Acids Res. 47, D419–D426 (2019).

33. Tsumagari, K. et al. Gene expression during normal and FSHD myogenesis. BMC Med. Genomics 4, (2011).

34. Banerji, C. R. S. et al. β-catenin is central to DUX4-driven network rewiring in facioscapulohumeral muscular dystrophy. J. R. Soc. Interface 12, (2015).

35. Lek, A. et al. Applying genome-wide CRISPR-Cas9 screens for therapeutic discovery in facioscapulohumeral muscular dystrophy. Sci. Transl. Med. 12, 9–11 (2020).

36. Grün, D. et al. De Novo Prediction of Stem Cell Identity using Single-Cell Transcriptome Data. Cell Stem Cell 19, 266–277 (2016).

37. Shannon, C. E. A Mathematical Theory of Communication. Bell Syst. Tech. J. 27, 623–656 (1948).

38. Macarthur, B. D. & Lemischka, I. R. Statistical mechanics of pluripotency. Cell 154, 484–489 (2013).

39. Banerji, C. R. S. et al. Cellular network entropy as the energy potential in Waddington's differentiation landscape. Sci. Rep. 3, 25–27 (2013).

40. Herman, J. S., Sagar & Grün, D. FateID infers cell fate bias in multipotent progenitors from single-cell RNA-seq data. Nat. Methods 15, 379–386 (2018).

41. Snel, B., Lehmann, G., Bork, P. & Huynen, M. A. String: A web-server to retrieve and display the repeatedly occurring neighbourhood of a gene. Nucleic Acids Res. 28, 3442–3444 (2000).

42. Szklarczyk, D. et al. STRING v11: Protein-protein association networks with increased coverage, supporting functional discovery in genome-wide experimental datasets. Nucleic Acids Res. 47, D607–D613 (2019).

43. Muraro, M. J. et al. A Single-Cell Transcriptome Atlas of the Human Pancreas. Cell Syst. 3, 385–394.e3 (2016).

44. Anders, S. et al. Differential expression analysis for sequence count data via mixtures of negative binomials. Adv. Environ. Biol. 7, 2803–2809 (2010).

